# *E*-scape: consumer specific landscapes of energetic resources derived from stable isotope analysis and remote sensing

**DOI:** 10.1101/2020.08.03.234781

**Authors:** W. Ryan James, Rolando O. Santos, Jennifer S. Rehage, Jennifer C. Doerr, James A. Nelson

## Abstract

Energy and habitat distribution are inherently linked. Energy is a major driver of the distribution of consumers, but estimating how much specific habitats contribute to the energetic needs of a consumer can be problematic. We present a new approach that combines remote sensing information and stable isotope ecology to produce maps of energetic resources (*E*-scapes). *E*-scapes project species specific resource use information onto the landscape to classify areas based on energetic importance and successfully predict the biomass and energy density of a consumer in salt marsh habitats in coastal Louisiana, USA. Our *E*-scape maps can be used alone or in combination with existing models to improve habitat management and restoration practices and have potential to be used to test fundamental movement theory.

## Introduction

The availability of energetic resources and habitat distribution are inherently linked. Habitats produce specific resources that are available to consumers, and energy is a major driver of consumer production, movement, and distribution (Wallace *et al.* 1999; Ware & Thomson 2005; Pyke 2019). The distribution of habitats, and therefore energy, is heterogeneous, and there is a substantial body of theoretical and empirical work that demonstrates how organisms respond to patterns of habitat and energy across landscapes (Wright 1983; Currie 1991; Guégan *et al.* 1998; Brown *et al.* 2004; Stein *et al.* 2014; Pyke 2019). This framework provides a link for how consumers are influenced by the distribution of energy and, coupled with technological advances in remote sensing and geographical information systems (GIS), provide an exciting opportunity to answer critical questions in spatial ecology and influence how we manage and restore rapidly changing ecosystems (Merkle *et al.* 2015; Fryxell *et al.* 2020).

An accurate species specific representation of resource availability at the landscape scale is required to test theories linking energy availability and species foraging or distribution. Spatial primary production estimates (e.g. normalized difference vegetation index (NDVI), chlorophyll-a concentration) and prey habitat suitability models are some of the approaches used to map resource availability for consumers across landscape and regional spatial scales (i.e., from 10s to 100s of kilometers) (Mosser *et al.* 2014; Abrahms *et al.* 2019; Geary *et al.* 2020). For example, a habitat suitability model of the dominant prey of brown pelicans (which included chlorophyll-a concentration as a model parameter) was used to test how foraging behavior changed during the breeding season (Geary *et al.* 2020). Landscape resource maps have typically focused on a single resource or prey species, which is accurate when a consumer specializes on that resource.

However, in many cases, a consumer is integrating multiple resources from different habitat types across the landscape. When a consumer is using multiple resources, mapping energy distribution is more difficult because resources are not produced evenly amongst habitats and consumers typically do not use all resources equally. Thus, in order to accurately represent energy distribution, information is needed on where resources are being produced across the landscape and the proportion each resource used by the consumer.

Remote sensing has long been used to produce landscape-level imagery of habitats, and digital platforms provide access and availability of satellite and aerial imagery more than ever before (Xie *et al.* 2008). Satellite programs like Landsat and Sentinel provide free multispectral imagery of the globe, and commercial satellites and unmanned aircraft systems (UAS) are becoming more affordable for providing high-resolution imagery (Tucker *et al.* 2004; Irons *et al.* 2012; Harris *et al.* 2019). GIS software can easily convert remotely sensed imagery into habitat cover maps, and remote sensing has helped in the mapping of different systems across multiple spatiotemporal scales. These new remote sensing products/maps can be combined with other spatially explicit data such as biogeochemical tracers, population information, or physical parameters to generate novel data products that can answer a wide array of ecological, management, and conservation questions (West *et al.* 2007; Effati *et al.* 2012).

Stable isotope ratios, typically of ^13^C/^12^C, ^15^N/^14^N, and ^34^S/^32^S, have been used for decades to determine the relative contributions of primary production sources in food webs (Peterson & Fry 1987; Fry 2007; Nelson *et al.* 2015). The general principle hinges literally upon the age-old adage “you are what you eat”. Organisms consume food and rearrange the consumed material to create new tissue. The stable isotope values, typically defined in del notation and expressed in per mil, of primary producers are controlled by a number of physical and biological processes that impart characteristic isotope values (Chanton *et al.* 1987; Farquhar *et al.* 1989). These characteristic values can then be traced as they are assimilated in the food web using Bayesian stable isotope mixing models (Stock *et al.* 2018). All plants fix carbon from the same atmospheric reservoir of CO_2_, currently −8 ‰ δ^13^C. For example, in coastal ecosystems carbon stable isotope values can be most useful in differentiating between C3 plants, such as mangroves, which fix carbon with a net fractionation of about −20 ‰ relative to the atmosphere and C4 plants, such as tropical and temperate salt tolerant grasses, which have a net fractionation of about −5 ‰ (Fry 2007). In the same systems sulfate reduction in sediments has large fractionation factor (30-70 ‰) and can be used as a strong indicator of pelagic vs. benthic primary production (Chanton *et al.* 1987; Nelson *et al.* 2012).

Here we present a method that combines stable isotope analysis, Bayesian mixing models, and remote sensing to build a landscape of energetic resources, or *E*-scape, for white shrimp (*Litopenaeus setiferus*) in Port Fourchon, LA. An *E*-scape combines the spatial locations where resources are being produced (habitat cover map) and how much of each resource the consumer is using (stable isotope analysis) to generate a species specific map of areas that contain habitats producing the resources being used by that species. Using our *E*-scapes, we investigated the relationship between energy distribution and white shrimp distribution and how the scale used to generate the *E*-scape mediated this relationship.

## Methods

Samples of white shrimp (*Litopenaeus setiferus*) were collected using a 1-m^2^ drop sampler at 55 randomly selected sampling locations in Port Fourchon, LA (Figure 1A) (Zimmerman *et al.* 1984; Nelson *et al.* 2019). We collected all of the white shrimp within the drop sampler to determine the abundance and biomass at each sampling location. Samples for stable isotope analysis and bomb calorimetry were removed, placed on ice, and frozen upon returning to the laboratory.

**Figure 1.**
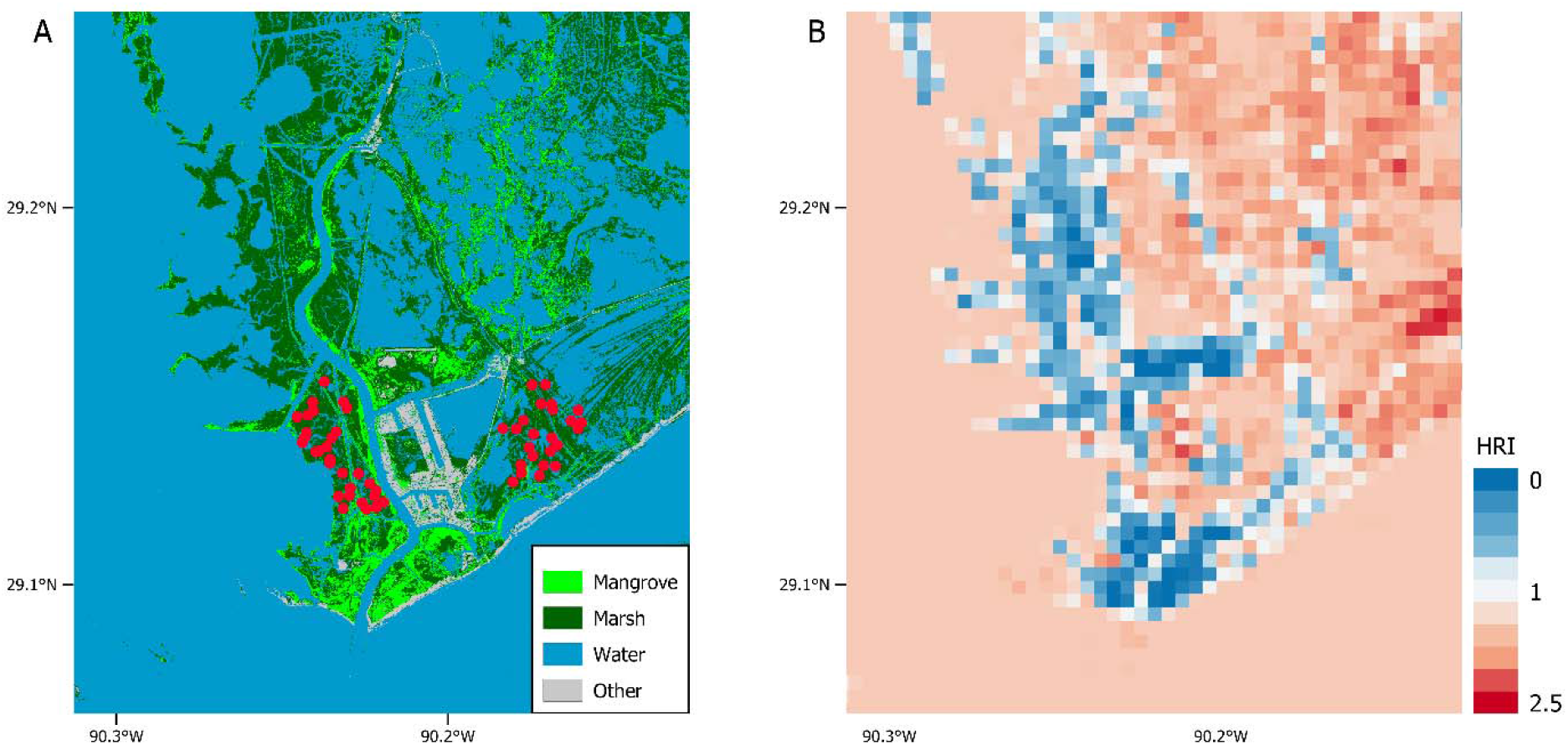
The Port Fourchon, LA A) habitat cover map showing the sampling locations of white shrimp (red points) and B) the corresponding white shrimp *E*-scape map. Warmer colors (HRI valuess > 1) are better energetically for white shrimp, and cooler colors (HRI values < 1) are worse energetically. The *E*-scape was generated at a cell size of 400 m × 400 m (similar area to a 200 m circle)

Primary production source and animal tissue samples were frozen at − 20°C in the laboratory until they could be processed for isotope analysis and bomb calorimetry. At each location, 5 individuals were pooled to create one composite sample. Samples were dried at 50◻C for 48 hours and ground. We determined the energy density (calories/g) of each sample using a Parr 6725 bomb calorimeter (Parr Instrument Company, Moline, IL, USA). We shipped samples to the Washington State University Stable Isotope Core Facility for C, N, and S content and stable isotope analysis. Carbon, nitrogen, and sulfur isotope values were calculated using the standard formula (Fry 2007). PeeDee Belemnite (PDB), atmospheric nitrogen, and Canyon Diablo Troilite (CDT) were used as the reference standards for C, N, and S, respectively. No C:N ratio was above 3.5; therefore, no lipid correction was applied (Layman *et al.* 2007; Nelson *et al.* 2011).

Bayesian mixing models were run in R using the package MixSIAR (Stock *et al.* 2018) to determine the relative basal resource contributions to shrimp at each sampling location. Each model was run with a Markov chain Monte Carlo algorithm that consisted of three chains, chain length of 3,000,000, burn-in of 1,500,000, and thin of 500 to ensure model convergence. Corrections were made for the elemental concentration in each source, and the trophic enrichment for each element was C = 1.0 ± 0.63 (mean ± sd), N = 3.0 ± 0.74, and S = 0.5 ± 0.2 (Phillips *et al.* 2014).

The *E*-scape of Port Fourchon, LA for white shrimp was made using the methods outlined in Figure 2. High-resolution aerial imagery from https://atlas.ga.lsu.edu was used to generate a habitat cover map of Port Fourchon, LA using the ‘Maximum Likelihood Classification’ tool in ArcGIS (v 10.5). This tool uses supervised classification maximum likelihood to assign a habitat class to each pixel of the image based on mean and variances of the habitat classes of the training data set. Four habitat classes were used: water, marsh, mangrove, and other. The ‘marsh’ class was comprised mainly of *Spartina alterniflora*, the ‘mangrove’ class was comprised mainly of *Avicennia germinans*, and the ‘other’ class was comprised mainly beach area and port facilities. Habitat cover areas were calculated using buffers with circle radius lengths of 50, 75, 100, 150, 200, 250, 300, 400, 500, 750, 1000, and 1500 m around the collection locations using the ‘landscapemetrics’ packages in R (Hesselbarth *et al.* 2019). White shrimp have a home range similar to that of the area of a 200 m radius circle (Rozas & Minello 1997; Webb & Kneib 2004; Nelson *et al.* 2019). The other size buffers were chosen to test the sensitivity of the *E*-scape at different scales. Edge habitat was calculated by measuring the linear distance between the water and vegetation (marsh and mangrove) habitat cover classes and multiplying by 2 m to generate an area. Edge area was calculated this way because benthic algae production is highest at the marsh edge (Wainright *et al.* 2000; Litvin *et al.* 2018), and benthic microalgae have recently been shown to have similar biomass at the edge habitat of both marsh and mangrove vegetation (Walker *et al.* 2019).

**Figure 2.**
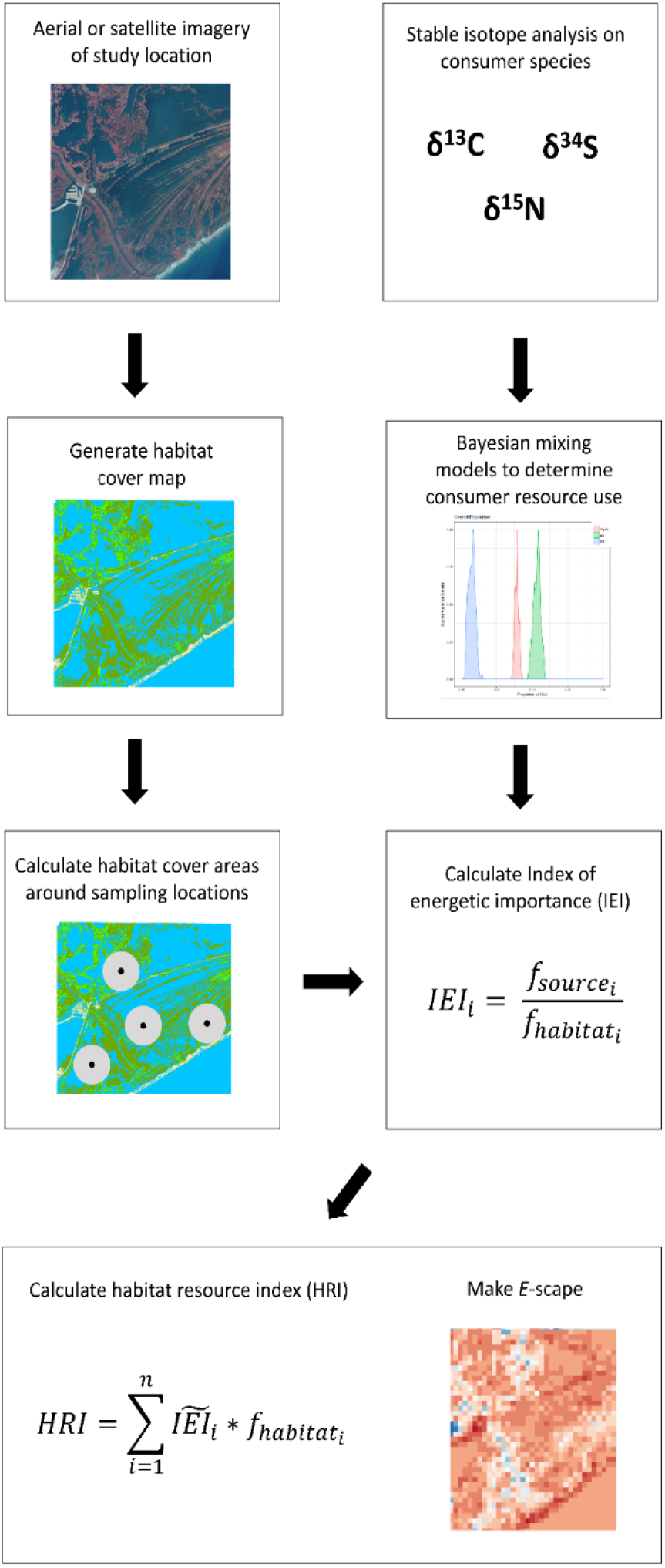
General methods for generating an *E*-scape

Habitat cover areas were combined with consumer resource use to calculate the index of energetic importance (IEI) for each basal resource and habitat type combination. Each IEI was calculated with the following formula:

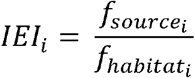

where 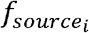 is the fraction of the contribution of source *i* to the total source use based on the results of the mixing model and 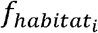 is the fraction of habitat *i* that produces source *i* to the overall area within the movement range of the consumer (area of the circle around the sampling point). An example of resource/habitat combination is amount of *Spartina alterniflora* derived production and the cover area of *S. alterniflora* marsh habitat. IEI values were calculated for phytoplankton/water, *Spartina*/marsh, and benthic algae/edge source/habitat combinations. The mangrove habitat source combination was not used in the analysis because resource use of mangrove was < 0.01. Each IEI is a measurement of how much energy of a resource a consumer is derived from relative to the amount of habitat that produces that resource where the consumer is foraging. An IEI around one means that the consumer is using a resource 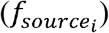 around the same amount as the proportion of the habitat that produces that resource relative to total area where that consumer is foraging over. An IEI greater than one means that the consumer is using that source more than expected based on the proportion of that habitat in the total foraging area, while the opposite is true for an IEI below one.

IEI values were combined with habitat cover areas to calculate the habitat resource index (HRI). HRI was calculated with the following formula:

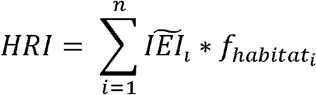

where 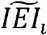 is the median of the IEI for the source/habitat combination *i* and 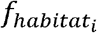 is the fraction of habitat *i* to the overall area within the movement range of the consumer. HRI is an index that represents a relative measurement of the quality of the habitats for producing the resources used by the consumer based on stable isotope analysis. An HRI value of 1 means that the area is producing the average amount of resources for the consumer. HRI values > 1 mean that the area is better for producing resources (i.e. more energy) for the consumer and the opposite is true for HRI values < 1 (Figure 1). The minimum possible HRI = 0, and the theoretical maximum for HRI is infinity, although it is very unlikely that this value will occur in nature because 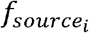 and 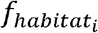 range between 0-1. Therefore, a unit of change is not linear for HRI, and log(HRI) should be used for linear modeling purposes so that unit change is the similar throughout the possible range of values.

One HRI value was calculated for each sampling location for the area enclosed within a circular buffer with the equation above. HRIs were calculated within a circular buffer with a radius length of 200 m based on field movement ranges of white shrimp in the field (Rozas & Minello 1997; Webb & Kneib 2004; Nelson *et al.* 2019). HRI values were also calculated at 50, 75, 100, 150, 250, 300, 400, 500, 750, 1000, and 1500 m radius circles around the sample points to test for the effect of scale. The HRI values were calculated using the mean IEIs that were calculated at the same scale (i.e. the IEIs calculated at 100 m were used in the calculation of the HRI at 100 m). A GLM with a gaussian error was used to test the relationship between log(HRI) and energy density (cal/g). GLMs with a gamma error and log link function were used to test the relationship between HRI and biomass, abundance, total calories (cal/g * biomass), and mean size (biomass/abundance). For each GLM outliers were removed if the value was outside of 1.5 ± the interquartile range. All analyses were done in R (R Core Team 2020).

## Results

White shrimp used benthic algae more than any other source (mean ± sd; 0.49 ± 0.04), followed by phytoplankton (0.38 ± 0.07), and *Spartina* (0.13 ± 0.04; Figure 3). Mangroves had a source contribution of < 0.01 of white shrimp (Figure 3).

**Figure 3.**
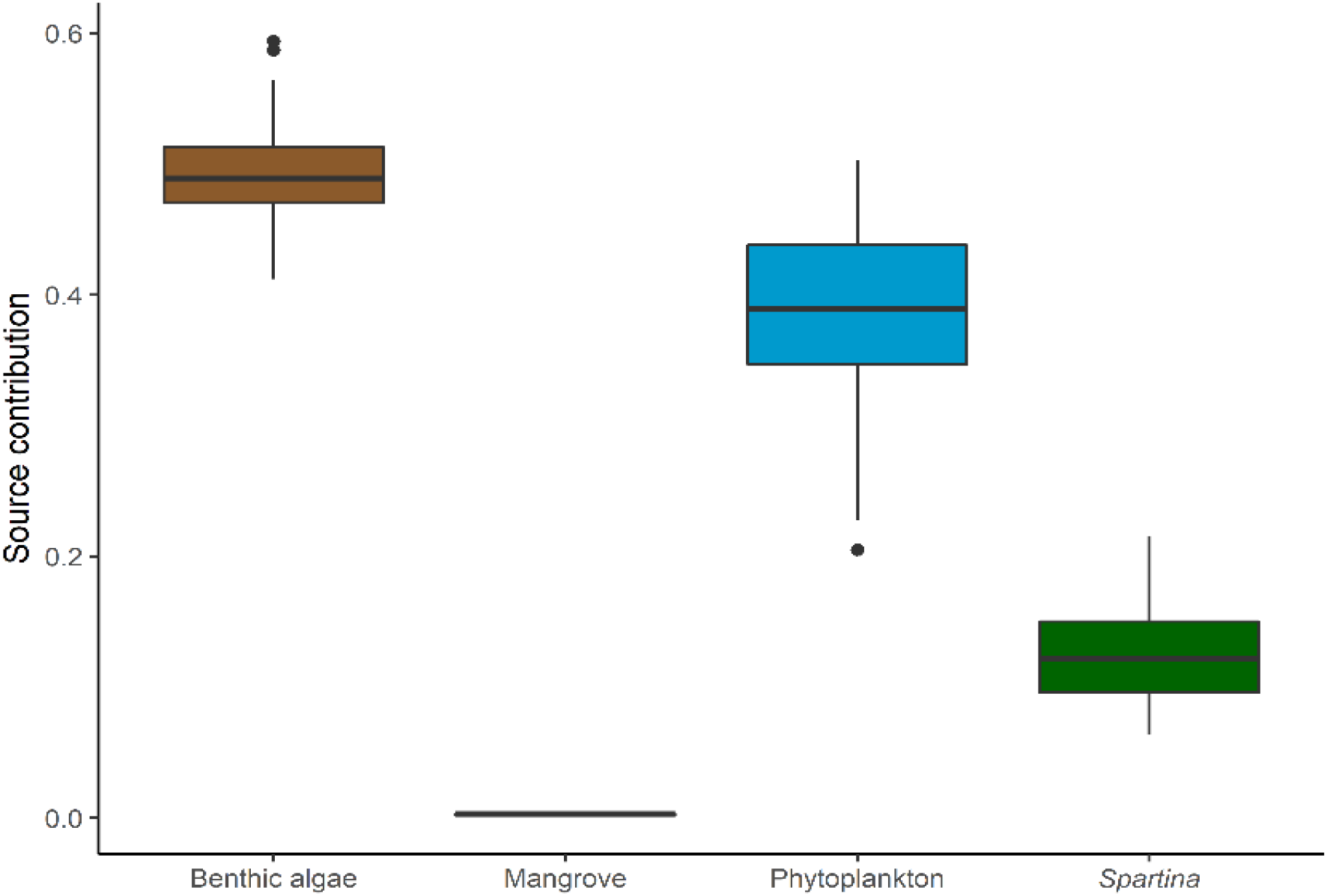
Bayesian mixing model results for white shrimp in Port Fourchon, LA.

The index of energetic importance (IEI) values are a representation of how much the white shrimp are using a resource relative to the amount of habitat that produces that resource (Table 1). Edge had consistently the highest IEI across all scales, with much smaller IEI values for both water and marsh (Table 1). Edge IEI values were highest at the smallest scale and declined until the 300 m radius, the lowest IEI value, where it increased as scale increased. Water IEI values were highest at the smallest scale and decreased as scale increase. Marsh IEI values were lowest at all scales of the three habitats and increased in value as scale increased.

**Table 1.** The index of energetic importance (IEI) values and interquartile ranges (IQR) for each source/habitat combination: benthic algae/edge, phytoplankton/water, and *Spartina*/marsh and the habitat resource index (HRI) values (mean ± SD) at varying scales of consumer foraging (size circle calculated around sampling location). HRI values > 1 are better than average energetically for white shrimp, while the opposite is true for HRI values < 1.

| Size | Edge IEI (IQR) | Water IEI (IQR) | Marsh IEI (IQR) | HRI (mean ± SD) |
| --- | --- | --- | --- | --- |
| 50 | 11.27 (6.61-28.12) | 3.02 (1.03-14.35) | 0.18 (0.14-0.25) | 1.38 ± 0.90 |
| 75 | 11.00 (6.91-14.13) | 1.72 (1.04- 4.71) | 0.19 (0.15-0.26) | 1.16 ± 0.57 |
| 100 | 9.19 (6.54-14.38) | 1.48 (1.01- 2.72) | 0.20 (0.15-0.27) | 1.07 ± 0.42 |
| 150 | 8.50 (6.66-12.97) | 1.36 (0.89- 1.87) | 0.21 (0.17-0.27) | 1.07 ± 0.36 |
| 200 | 8.19 (6.72-11.26) | 1.26 (0.98- 1.75) | 0.21 (0.18-0.26) | 1.04 ± 0.32 |
| 250 | 8.28 (6.48-10.79) | 1.29 (0.98- 1.69) | 0.22 (0.18-0.25) | 1.07 ± 0.30 |
| 300 | 8.16 (6.58-10.49) | 1.20 (0.92- 1.52) | 0.21 (0.18-0.26) | 1.04 ± 0.26 |
| 400 | 8.38 (6.95-10.36) | 1.14 (0.90- 1.37) | 0.22 (0.19-0.27) | 1.03 ± 0.22 |
| 500 | 8.42 (7.02-10.89) | 1.10 (0.85- 1.34) | 0.24 (0.18-0.27) | 1.02 ± 0.20 |
| 750 | 9.16 (7.29-11.34) | 0.96 (0.78- 1.12) | 0.25 (0.18-0.30) | 1.02 ± 0.14 |
| 1000 | 9.38 (7.84-11.54) | 0.87 (0.74- 1.06) | 0.27 (0.20-0.34) | 0.99 ± 0.11 |
| 1500 | 9.92 (8.33-11.87) | 0.79 (0.69- 0.98) | 0.31 (0.23-0.38) | 0.99 ± 0.11 |

Habitat resource index (HRI) values at the 200 m scale were 1.04 ± 0.32 (mean ± SD) around the sampling locations (Table 1). HRI values are a relative metric of quality of the habitats for producing resources used by the white shrimp and were highest in areas that contained the most edge habitat (Figure 1). There was a significant relationship between HRI value and body size (*t*-value = 4.8, p < 0.001), abundance (*t*-value = 2.5, p = 0.018), biomass (*t*-value = 5.4, p < 0.001), and total calories (*t*-value = 5.1, p < 0.001) at the 200 m scale (Figure 4, Table S1). The relationship between HRI values and energy density (calories/g) was not significant (p > 0.05). For the other scales, the relationship between HRI values and body size was significant (p < 0.05) at intermediate scales (100 – 750 m, Table S1). At the 150-250 m scales, there was a significant relationship with HRI values and abundance (Table S1). There was a signification relationship (p < 0.05) between HRI value and biomass for all but the 1500 m scale. The same was true for total calories (Table S1). There was no significant relationship between HRI value and Calories/g at any scale.

**Figure 4.**
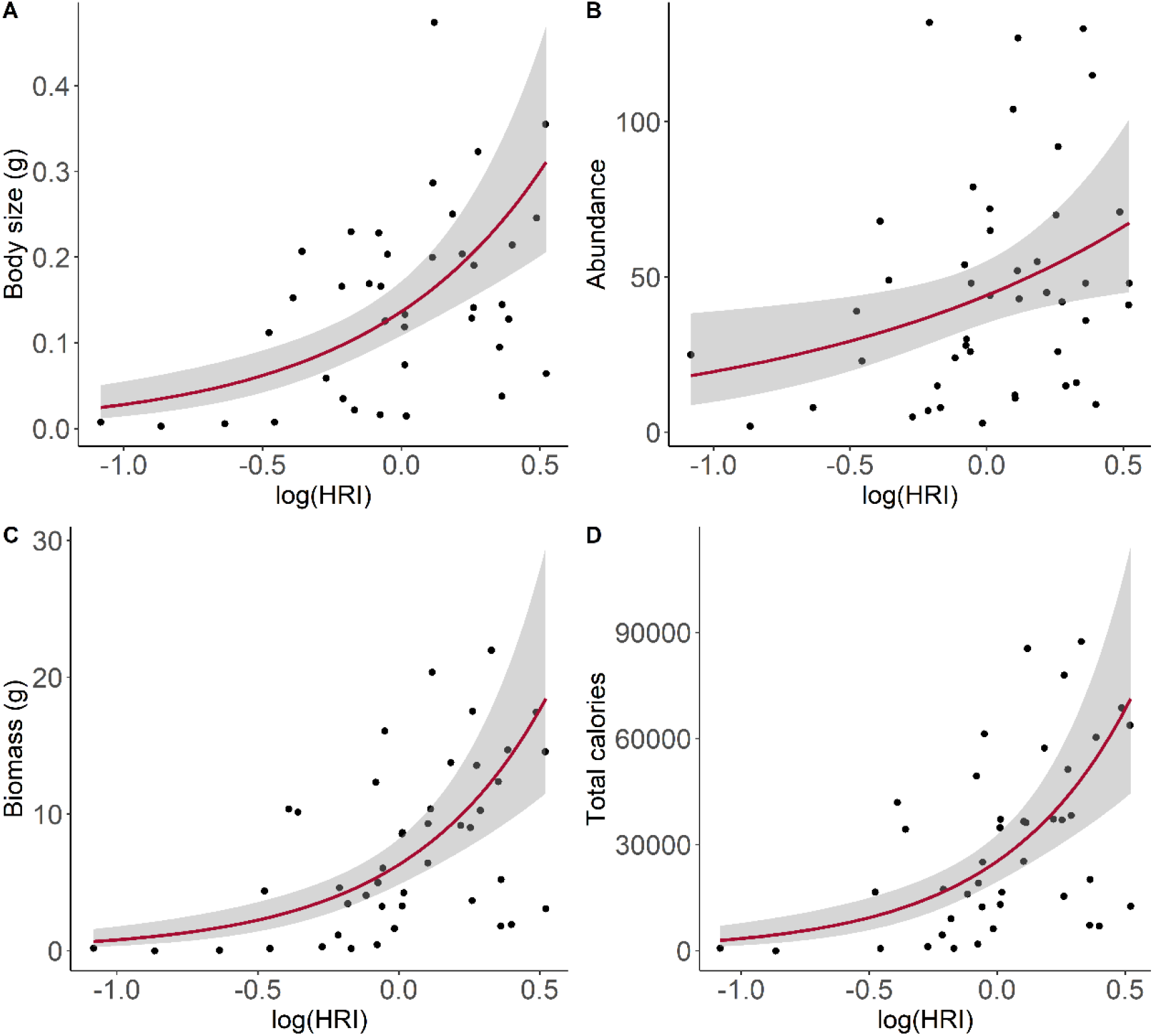
The relationship between habitat resource index and white shrimp A) body size, B) abundance, C) biomass, and D) total calories. HRI values were calculated within a 200 m radius circle around sampling locations.

## Discussion

Our results demonstrate that *E*-scapes can predict the spatial distribution of biomass and energetic density of a consumer by combing spatial habitat and resource use data (Figure 4). White shrimp size, abundance, biomass, and total calories increased as the habitat resource index increased across the marsh seascape (Figure 4). Individual white shrimp energy density (cal/g) was not related to energy distribution. These results are supported by previous work that showed white shrimp energy density did not change depending on the habitat type of the shrimp (Nelson *et al.* 2019).

Habitat resource index (HRI) values predicted white shrimp distribution within its foraging range (200 m), but not at all scales tested. At scales less than 200 m the areas sampled failed to include all the habitats and resources used by shrimp creating an oversampling artifact. At the larger scales, the opposite is true, and the forage areas were over aggregated leading to poor representation of foraging habitat. These results demonstrate that choosing the right scale for generating the *E*-scape is important and should correspond to the foraging range of the consumer. For example, consumers that are foraging over much larger areas than shrimp (e.g. whale or bird), would require a larger *E*-scape sampling unit on the order kilometers instead of meters (Abrahms *et al.* 2019; Geary *et al.* 2020). New tracking techniques can be used to inform these scales which were previously poorly understood (Abrahms *et al.* 2019; Geary *et al.* 2020).

The index of energetic importance (IEI) represents how much a consumer is using a resource relative to the amount of habitat that is producing that resource. White shrimp are derived of 49% benthic algae and 38% phytoplankton, but since there is much less edge habitat (the habitat where benthic algae is produced), the IEI for edge is almost an order of magnitude larger than the IEI for water (Table 1). Therefore, the habitats that contain the most edge habitat are of the highest energetic importance for white shrimp (Figure 1). The IEI for marsh is < 1 at all scales indicating that white shrimp use energetic resources from the marsh at a lower rate than their availability in the system (Table 1). Although areas that contain a high amount of marsh habitat are less favorable energetically than the average habitat (HRI < 1), these habitats are still producing energy being used by white shrimp and are more energetically favorable than areas of high mangrove habitat (which white shrimp are not using as an energy source, Figure 3,(Nelson *et al.* 2019). Thus, the maps can differentiate between habitats suitable to occupy vs habitats that are producing energy.

In our calculation of HRI and IEI values, the fraction of habitat 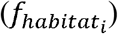 is based on the area of habitat cover. This calculation assumes that all areas of a given habitat type have an equal chance of producing a resource. For example, we make the assumption that all areas of water in our habitat cover map (Figure 1A) have an equal chance of producing phytoplankton. This assumption may not be acceptable in all applications, especially when applying these methods to consumers that have very large foraging ranges (Geary *et al.* 2020). For these cases, modifications can be made to 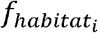 to incorporate the spatial differences in production such as incorporating chlorophyll-a maps or lidar data to incorporate the three dimensional structure of the habitats. One limitation to our approach is that phytoplankton is produced in three dimensions, unlike the other sources, and we are presently not able to account for the three-dimensional structure of water across the seascape with the available data. Accounting for water volume will be especially important in systems that are stratified or in which phytoplankton production is integrated over a significant depth (Cole & Cloern 1984). One way to incorporate volume into 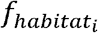 is to modify by accounting for the depth of the habitat in relation to the euphotic zone of the system (Cole & Cloern 1984). Unfortunately, this type of data is not always available and was not available in our study area. Other modifications could include parameters that include temporal differences in access to habitats which can be major drivers of foraging behaviors of consumers (Nelson *et al.* 2015).

These *E*-scape maps allow users to identify key areas of the landscape in terms of their importance to the energetic requirements of a consumer. Researchers could apply *E*-scape maps to conservation, management, or restoration questions to identify areas of importance and to take management action. In combination with other parameters, *E*-scape maps could improve habitat suitability models and integrate energetics into existing modelling frameworks. Similar approaches have been applied to terrestrial ecosystems to investigate population and movement responses of large-bodied herbivores (Merkle *et al.* 2015; Fryxell *et al.* 2020). For example, the population viability of caribou was determined by modeling the response to resource distribution as well as other environmental and biological factors (Fryxell *et al.* 2020). Field observations of diet and grazing amount to determine digestible energy content and combined with habitat cover maps were used quantify the distribution of energy (Fryxell *et al.* 2020). Although effective, this technique requires extensive field work and data, and is limited to terrestrial herbivores where the direct measurements of grazing can occur. Our method improves upon previous methods by using stable isotope analysis, which provides a representation of the assimilated energy for which a consumer is derived (Layman *et al.* 2012). With stable isotope analysis and Bayesian mixing models, estimates of consumer resource use are not limited to consumers where direct consumption can be observed (e.g. terrestrial herbivores), expanding the number of ecosystems and types of consumers that can studied.

Our study links energy to population and energetic distribution of white shrimp, but if paired with tracking data *E*-scapes have the capability to further our understanding of consumer movement and foraging. Optimal Foraging Theory predicts that consumers will optimize net energy intake per unit time foraging and consumers would be expected to spend more time foraging in areas of greater resources (MacArthur & Pianka 1966). Therefore, *E*-scape maps describe a “null model” to test Optimal Foraging Theory for a particular consumer. Tracking data can be used in combination of *E*-scapes to test foraging strategies in the context of energy distribution (e.g. even vs patchy distribution) or paired with other spatial environmental (e.g. salinity, temperature) or biotic factors (e.g. predation risk) to identify key drivers of movement and test hypotheses on variations of OFT. Recent studies have focused on consumers optimizing foraging by tracking temporal resource waves but have been limited to systems with discrete waves of a dominant energy source (Mosser *et al.* 2014; Abrahms *et al.* 2019). Because our approach quantifies which energy sources a consumer is using, it is an improvement of mapping energy distribution. *E*-scapes will expand the systems where foraging patterns can be tested in the field, especially when resources do not have discrete waves and spatial and spatiotemporal variation dominate where resources are located, expanding our understanding of consumer foraging. *E*-scapes can be used alone or in combination with existing models to test fundamental movement theory and improve habitat management and restoration practices.

## Acknowledgements

We thank Laura McDonald, Holly Mayeux, and Victoria Furka for assistance processing samples in the laboratory. Justin Lesser, David Behringer, Juan Salas, Lawrence Rozas, and Shawn Hillen participated in the field collections. Benjamin Harlow was responsible for running the stable isotope analysis. This work was supported by the National Oceanic and Atmospheric Administration, National Marine Fisheries Service, Louisiana Sea Grant, The National Academies of Science, Engineering, and Medicine Gulf Research Program, and NSF (DEB-1832229).

## Authorship

WRJ, ROS, JSR and JAN designed the study. WRJ and JD collected and processed samples. WRJ analyzed the data. WRJ wrote the first draft with input from JAN. All authors contributed substantially to revising the manuscript.

## Data Accessibility

Data will be archived in an appropriate public repository and the data DOI will be included at the end of the article.

## Supporting information

Model results for GLMs testing the relationship between the log(HRI) and different biomass and energy measurements of for white shrimp at each scale tested. Significant (p < 0.05) model results in bold. T = *t*-vale, AIC = Akaike information criterion

| Size | Size |  |  | Abundance |  |  | Biomass |  |  | Calories/g |  |  | Total calories |  |  |
| --- | --- | --- | --- | --- | --- | --- | --- | --- | --- | --- | --- | --- | --- | --- | --- |
|  | T | P value | AIC | T | P value | AIC | T | P value | AIC | T | P value | AIC | T | P value | AIC |
| 50 | 1.0 | 0.317 | -68.8 | 0.3 | 0.780 | 436.8 | **2.7** | **0.009** | **265.6** | 0.3 | 0.733 | 698.2 | **2.6** | **0.013** | **951.6** |
| 75 | 1.5 | 0.132 | -69.6 | 1.2 | 0.219 | 435.6 | **2.6** | **0.012** | **265.4** | 0.6 | 0.545 | 697.9 | **2.3** | **0.024** | **951.9** |
| 100 | **1.8** | **0.078** | **-70.5** | 1.4 | 0.158 | 435.1 | **2.2** | **0.031** | **266.1** | 1.1 | 0.292 | 697.2 | **2.0** | **0.049** | **952.3** |
| 150 | **4.1** | **0.000** | **-76.1** | **2.4** | **0.019** | **432.5** | **4.5** | **0.000** | **260.0** | 0.5 | 0.627 | 698.1 | **4.3** | **0.000** | **946.8** |
| 200 | **4.8** | **0.000** | **-78.6** | **2.5** | **0.018** | **432.3** | **5.4** | **0.000** | **256.9** | 0.6 | 0.555 | 698.0 | **5.1** | **0.000** | **943.7** |
| 250 | **4.2** | **0.000** | **-76.9** | **2.0** | **0.049** | **433.6** | **5.0** | **0.000** | **258.0** | 0.8 | 0.413 | 697.6 | **4.9** | **0.000** | **944.3** |
| 300 | **3.2** | **0.003** | **-73.9** | 1.6 | 0.126 | 434.9 | **4.1** | **0.000** | **261.5** | 1.0 | 0.324 | 697.3 | **4.0** | **0.000** | **947.3** |
| 400 | **2.8** | **0.007** | **-72.9** | 1.2 | 0.232 | 435.7 | **3.8** | **0.001** | **262.9** | 0.8 | 0.408 | 697.6 | **3.7** | **0.001** | **948.6** |
| 500 | **3.0** | **0.005** | **-72.9** | 0.9 | 0.382 | 436.2 | **3.8** | **0.000** | **262.8** | 0.9 | 0.389 | 697.5 | **3.8** | **0.001** | **948.2** |
| 750 | **2.4** | **0.020** | **-71.1** | 0.0 | 0.995 | 436.9 | **3.0** | **0.004** | **265.3** | 0.8 | 0.453 | 697.7 | **2.9** | **0.006** | **950.9** |
| 1000 | 2.0 | 0.054 | -70.0 | -0.8 | 0.416 | 436.3 | **2.2** | **0.037** | **267.5** | 1.1 | 0.267 | 697.0 | **2.1** | **0.046** | **953.2** |
| 1500 | 1.2 | 0.225 | -68.9 | -1.1 | 0.277 | 435.9 | 1.0 | 0.302 | 269.0 | 1.4 | 0.154 | 696.2 | 1.0 | 0.303 | 954.6 |

## References

Abrahms, B., Hazen, E.L., Aikens, E.O., Savoca, M.S., Goldbogen, J.A., Bograd, S.J., et al. (2019). Memory and resource tracking drive blue whale migrations. Proc Natl Acad Sci USA, 116, 5582–5587.

Brown, J.H., Gillooly, J.F., Allen, A.P., Savage, V.M. & West, G.B. (2004). Toward a metabolic theory of ecology. Ecology, 85, 1771–1789.

Chanton, J.P., Martens, C.S. & Goldhaber, M.B. (1987). Biogeochemical cycling in an organic-rich coastal marine basin. 8. A sulfur isotopic budget balanced by differential diffusion across the sediment-water interface. Geochimica et Cosmochimica Acta, 51, 1201–1208.

Cole, B. & Cloern, J. (1984). Significance of biomass and light availability to phytoplankton productivity in San Francisco Bay. Mar. Ecol. Prog. Ser., 17, 15–24.

Currie, D.J. (1991). Energy and large-scale patterns of animal-and plant-species richness. The American Naturalist, 137, 27–49.

Effati, M., Rajabi, M.A., Samadzadegan, F. & Blais, J.R. (2012). Developing a novel method for road hazardous segment identification based on fuzzy reasoning and GIS. Journal of Transportation Technologies, 2, 32.

Farquhar, G.D., Ehleringer, J.R. & Hubick, K.T. (1989). Carbon isotope discrimination and photosynthesis. Annual review of plant biology, 40, 503–537.

Fry, B. (2007). Stable isotope ecology. Springer Science & Business Media.

Fryxell, J.M., Avgar, T., Liu, B., Baker, J.A., Rodgers, A.R., Shuter, J., et al. (2020). Anthropogenic Disturbance and Population Viability of Woodland Caribou in Ontario. The Journal of Wildlife Management, 84, 636–650.

Geary, B., Leberg, P.L., Purcell, K.M., Walter, S.T. & Karubian, J. (2020). Breeding Brown Pelicans Improve Foraging Performance as Energetic Needs Rise. Scientific Reports, 10, 1686.

Guégan, J.-F., Lek, S. & Oberdorff, T. (1998). Energy availability and habitat heterogeneity predict global riverine fish diversity. Nature, 391, 382–384.

Harris, J.M., Nelson, J.A., Rieucau, G. & Broussard III, W.P. (2019). Use of Drones in Fishery Science. Transactions of the American Fisheries Society, 0.

Hesselbarth, M.H., Sciaini, M., With, K.A., Wiegand, K. & Nowosad, J. (2019). landscapemetrics: an open◻source R tool to calculate landscape metrics. Ecography, 42, 1648–1657.

Irons, J.R., Dwyer, J.L. & Barsi, J.A. (2012). The next Landsat satellite: The Landsat data continuity mission. Remote Sensing of Environment, 122, 11–21.

Layman, C.A., Araujo, M.S., Boucek, R., Hammerschlag◻Peyer, C.M., Harrison, E., Jud, Z.R., et al. (2012). Applying stable isotopes to examine food◻web structure: an overview of analytical tools. Biological Reviews, 87, 545–562.

Layman, C.A., Arrington, D.A., Montaña, C.G. & Post, D.M. (2007). Can stable isotope ratios provide for community-wide measures of trophic structure? Ecology, 88, 42–48.

Litvin, S.Y., Weinstein, M.P., Sheaves, M. & Nagelkerken, I. (2018). What Makes Nearshore Habitats Nurseries for Nekton? An Emerging View of the Nursery Role Hypothesis. Estuaries and Coasts, 1–12.

MacArthur, R.H. & Pianka, E.R. (1966). On optimal use of a patchy environment. The American Naturalist, 100, 603–609.

Merkle, J.A., Cherry, S.G. & Fortin, D. (2015). Bison distribution under conflicting foraging strategies: site fidelity vs. energy maximization. Ecology, 96, 1793–1801.

Mosser, A.A., Avgar, T., Brown, G.S., Walker, C.S. & Fryxell, J.M. (2014). Towards an energetic landscape: Broad◻scale accelerometry in woodland caribou. Journal of Animal Ecology, 83, 916–922.

Nelson, J., Chanton, J., Coleman, F. & Koenig, C. (2011). Patterns of stable carbon isotope turnover in gag, Mycteroperca microlepis, an economically important marine piscivore determined with a non-lethal surgical biopsy procedure. Environmental Biology of Fishes, 90, 243–252.

Nelson, J.A., Deegan, L. & Garritt, R. (2015). Drivers of spatial and temporal variability in estuarine food webs. Marine Ecology Progress Series, 533, 67–77.

Nelson, J.A., Lesser, J., James, W.R., Behringer, D.P., Furka, V. & Doerr, J.C. (2019). Food web response to foundation species change in a coastal ecosystem. Food Webs, 21, e00125.

Nelson, J.A., Wilson, R.M., Coleman, F.C., Koenig, C.C., DeVries, D., Gardner, C., et al. (2012). Flux by fin: fish mediated carbon and nutrient flux in the northeastern Gulf of Mexico. Marine Biology, 159, 365–372.

Peterson, B.J. & Fry, B. (1987). Stable isotopes in ecosystem studies. Annual review of ecology and systematics, 18, 293–320.

Phillips, D.L., Inger, R., Bearhop, S., Jackson, A.L., Moore, J.W., Parnell, A.C., et al. (2014). Best practices for use of stable isotope mixing models in food-web studies. Canadian Journal of Zoology, 92, 823–835.

Pyke, G. (2019). Animal movements: an optimal foraging approach. In: Encyclopedia of animal behavior. Elsevier Academic Press, pp. 149–156.

R Core Team. (2020). R: A Language and Environment for Statistical Computing. R Foundation for Statistical Computing, Vienna, Austria.

Rozas, L.P. & Minello, T.J. (1997). Estimating densities of small fishes and decapod crustaceans in shallow estuarine habitats: a review of sampling design with focus on gear selection. Estuaries, 20, 199–213.

Stein, A., Gerstner, K. & Kreft, H. (2014). Environmental heterogeneity as a universal driver of species richness across taxa, biomes and spatial scales. Ecology letters, 17, 866–880.

Stock, B.C., Jackson, A.L., Ward, E.J., Parnell, A.C., Phillips, D.L. & Semmens, B.X. (2018). Analyzing mixing systems using a new generation of Bayesian tracer mixing models. PeerJ, 6, e5096–e5096.

Tucker, C.J., Grant, D.M. & Dykstra, J.D. (2004). NASA’s global orthorectified Landsat data set. Photogrammetric Engineering & Remote Sensing, 70, 313–322.

Wainright, S., Weinstein, M., Able, K. & Currin, C. (2000). Relative importance of benthic microalgae, phytoplankton and the detritus of smooth cordgrass Spartina alterniflora and the common reed Phragmites australis to brackish-marsh food webs. Mar. Ecol. Prog. Ser., 200, 77–91.

Walker, J.E., Angelini, C., Safak, I., Altieri, A.H. & Osborne, T.Z. (2019). Effects of changing vegetation composition on community structure, ecosystem functioning, and predator-prey interactions at the Saltmarsh-Mangrove Ecotone. Diversity, 11, 208.

Wallace, J.B., Eggert, S.L., Meyer, J.L. & Webster, J.R. (1999). Effects of resource limitation on a detrital◻based ecosystem. Ecological monographs, 69, 409–442.

Ware, D.M. & Thomson, R.E. (2005). Ecology: Bottom-up ecosystem trophic dynamics determine fish production in the northeast pacific. Science, 308, 1280–1284.

Webb, S. & Kneib, R.T. (2004). Individual growth rates and movement of juvenile white shrimp (Litopenaeus setiferus) in a tidal marsh nursery. Fishery Bulletin, 102, 376–388.

West, J.B., Ehleringer, J.R. & Cerling, T.E. (2007). Geography and vintage predicted by a novel GIS model of wine δ18O. Journal of Agricultural and Food Chemistry, 55, 7075–7083.

Wright, D.H. (1983). Species-energy theory: an extension of species-area theory. Oikos, 496–506.

Xie, Y., Sha, Z. & Yu, M. (2008). Remote sensing imagery in vegetation mapping: a review. Journal of plant ecology, 1, 9–23.

Zimmerman, R.J., Minello, T.J. & Zamora, G. (1984). Selection of vegetated habitat by brown shrimp, Penaeus aztecus, in a Galveston Bay salt marsh. Fishery Bulletin, 82, 325–336.

